# Portable X-ray Fluorescence for Measuring Lead in the Hair of Wild Mammals

**DOI:** 10.1101/2023.03.13.531209

**Authors:** Evie M. Jones, Andrew J. Bengsen, Aaron J. Specht, Amelia J. Koch, Rodrigo K. Hamede, Menna E. Jones, Jordan O. Hampton

**Affiliations:** School of Natural Sciences, University of Tasmania, Private Bag 55, Hobart, TAS, 7001, Australia; Vertebrate Pest Research Unit, NSW Department of Primary Industries, 4 Marsden Park Rd, Calala, New South Wales 2340, Australia; Biosphere Environmental Consultants, Tamworth, NSW 2340, Australia; School of Health Sciences, Purdue University, 610 Purdue Mall, West Lafayette, Indiana 47907, United States; Forest Practices Authority, 30 Patrick St, Hobart, TAS, 7001, Australia; Faculty of Science, University of Melbourne, Parkville, Victoria 3052, Australia; Harry Butler Institute, Murdoch University, 90 South Street, Murdoch, Western Australia 6150, Australia

**Keywords:** ecotoxicology, marsupial, non-invasive, Pb, scavenger, Tasmanian devil

## Abstract

Lead exposure threatens scavenging wildlife globally. For inexpensive estimation of lead concentration in bones from avian scavengers, portable X-ray fluorescence (XRF) devices have been trialed. However, portable XRF has not been validated for tissue lead measurement in non-human mammalian scavengers. We evaluated portable XRF for hair lead measurement in the endangered Tasmanian devil (*Sarcophilus harrisii*). We first analyzed large (∼1.0 g) hair samples from 39 deceased animals from southeastern Tasmania via portable XRF and then inductively-coupled plasma mass spectrometry (ICP-MS) (validation study). We then measured lead concentrations via portable XRF only in 61 small (∼0.1 g) hair samples from live devils from a plantation landscape (plantation study). Portable XRF measures of hair lead levels were positively correlated with ICP-MS values (*R*^2^ = 0.61). In the validation study, 95% of sampled Tasmanian devils had relatively low lead levels (< 2 mg/kg), but the remaining two showed elevated lead levels (> 15 mg/kg). Mean lead levels in the plantation study and validation study were not significantly different. Our preliminary results suggest that portable XRF can provide a useful measure of differences in lead levels in wildife hair over a coarse scale. We provide recommendations for further research and refinement of this method.

**Synopsis:** Portable XRF can provide inexpensive and non-destructive analysis of environmental contaminants in wildlife. We present the first evaluation of portable XRF for analysis of lead contamination in mammalian scavenger hair.

## INTRODUCTION

Lead is one of the best-known toxicants, with deleterious effects on humans, animals and plants. Recent research has identified lead-based ammunition as a major source of anthropogenic environmental lead globally,^1,2^ representing a significant health hazard to wildlife^3^ and potentially humans^4^, leading to calls for a transition to non-toxic alternatives^5,6^. This source of lead is a particular threat to scavenging wildlife^7^. As a result of exposure to lead-based ammunition, numerous species of avian scavengers have been found with elevated tissue lead concentrations^3,8,9^. Impacts of lead exposure on avian scavengers include increased mortality^10^, suppressed population growth rates^11^, altered spatial behavior^12^ and inhibition of enzymes^13^, but little is known about the health impacts of lead exposure on wild mammalian scavengers. Lead exposure is a global issue, but relatively little research has been conducted in Australia^6^. The exceptions to this trend have been studies of Australia’s top avian predator, the wedge-tailed eagle (*Aquila audax*)^14–16^, and one marsupial scavenger, the Tasmanian devil (*Sarcophilus harrisii*)^17,18^. However, many more wildlife species may be at risk in Australia, given that wildlife shooting (culling, hunting and commercial harvesting) is ubiquitous in Australia, and relies almost exclusively on lead-based ammunition^6,19^.

While scavenging birds have been relatively well-studied, comparatively few studies have assessed the effects of lead pollution from ammunition on wild mammalian scavengers^20^. The notable exception to this trend has been studies of bears (*Ursus* spp.) in the Northern Hemisphere^21–24^. One of the few studies from the Southern Hemisphere demonstrated harmful blood lead levels in Tasmanian devils in captivity^25^. The Tasmanian devil is a morphologically specialized scavenger and the largest marsupial carnivore in the world^26^. They are endangered, with population declines of over 80% since 1996 caused by a transmissible cancer, devil facial tumor disease^27^. The low population size of the species and their reliance on scavenging makes the potential impact of lead contamination particularly concerning.

Due to their endangered status, the Tasmanian devil is the focus of intense conservation efforts, including proactive attempts to identify threatening processes that may prevent or delay their recovery, such as habitat modification and vehicle collisions^28–30^. There are concerns that lead exposure may also threaten the recovery of devils. The devil populations that are likely to be most affected by lead pollution are those inhabiting modified anthropogenic landscapes where culling of browsing mammals is common. For instance, in Tasmanian timber plantations (henceforth ‘plantations’), culling of marsupial browsers is regularly conducted prior to planting in coupes to protect seedlings^31^. The browsers targeted include brushtail possums (*Trichosurus vulpecula*), Bennett’s wallabies (*Notamacropus rufogriseus*), and rufous-bellied pademelons (*Thylogale billardierii*)^32^. All three species constitute primary prey for devils^33^, and currently culling methods involve the use of lead-based ammunition^18^.

The studies of Hivert et al. (2018)^17^ and Hutchinson et al. (2023)^18^ measured lead exposure in wild devils via blood and liver, respectively, and reported relatively low lead concentrations. However, these studies used samples from across Tasmania. No study has specifically investigated lead exposure in devils living in a landscape subjected to regular culling of browsers. To our knowledge, all past studies on lead exposure in scavenging mammals have used invasive sampling, whether samples were collected from live or dead animals. Tissue types analyzed have included blood^21,22,24,25,34^, teeth^23^ and liver^18,35^. However, recent reviews have highlighted the value of non-invasive sampling of heavy metal exposure in wild mammals^36^.

One non-invasive approach to investigating heavy metal exposure in mammals is through hair analysis, a technique used by some recent studies on scavengers such as brown bears (*Ursus arctos*)^37^ and non-scavenging species such as fruit bats (*Pteropus* spp.)^38^, Malay civets (*Viverra tangalunga*)^39^ and roe deer (*Capreolus capreolus*)^40^. Hair reflects an individual’s heavy metal exposure during the period of hair growth^41^. Collecting hair samples from wild animals can be done remotely using hair snares^42^ or during handling of captured wild animals^38^.

Most studies on lead exposure in mammalian scavengers have used *ex situ* analysis via inductively-coupled plasma mass spectrometry (ICP-MS) in a laboratory^22–24^, which is expensive and time-consuming. X-ray fluorescence (XRF) is another option for analyzing lead in tissues such as hair^43^, horn^44^ and bone^45,46^, and has long been used in human medicine^47^. The XRF device analyzes the elemental content of samples through the fluorescence produced by X-rays from a low energy x-ray tube (50 kV). XRF methods have been used to determine lead levels in the hair of humans^43,48,49^ but not, to our knowledge, other mammals.

Portable XRF units have recently become commercially available. As they are handheld and battery-powered, these units are relatively inexpensive, quick and practical for use in the field, with most functions taking only three minutes to measure samples^14^. Advantages of portable XRF over ICP-MS include lower cost and non-destructive analysis of samples^14,50^. One important disadvantage of XRF is that the instrument emits ionizing radiation with corresponding risks associated with health and safety for users^51^. Portable XRF has been validated for measurement of lead in bones of several avian species, showing strong correlations with ICP-MS values, such as wedge-tailed eagles^14^, California condors (*Gymnogyps californianus*)^46^, Pacific black ducks (*Anas superciliosa*)^52^, and common loons (*Gavia immer*)^45^. However, portable XRF has not been validated for measurement of lead in the hair of non-human mammals.

Here, we investigated hair lead concentrations in wild Tasmanian devils in two stages. The first stage was a validation study (comparing portable XRF to traditional ICP-MS methods) and the second stage was a study of lead exposure in a forestry plantation landscape using XRF. Our aims were: 1) to determine if portable XRF measures of hair lead levels in Tasmanian devils are similar enough to ICP-MS results that they can be used as a proxy, 2) to determine the usefulness of portable XRF as a tool to rapidly identify lead exposure levels in mammals through non-destructive analysis of hair samples, and 3) to use portable XRF to non-invasively identify lead exposure in an endangered mammalian scavenger living in a landscape subject to browser culling operations.

## MATERIALS AND METHODS

### Validation study

Hair from Tasmanian devils (n = 39) was collected from archived collections held in the Tasmanian Museum and Art Gallery (TMAG) and the University of Tasmania (UTAS). Road-killed devils were collected between April 2017 and February 2022 from across Tasmania, concentrated in the southeast, which is dominated by native forest and agricultural land (Figure 1). Approximately 1 g of hair was shaved from whole frozen devil carcasses and transported in ice packs to the laboratory for analysis. Lead concentrations in each hair sample were analyzed with both portable XRF and ICP-MS. As this study aimed in part to validate portable XRF for use in the field, we did not wash hair samples prior to analysis, as this would be impractical in the field.

**Figure 1.**
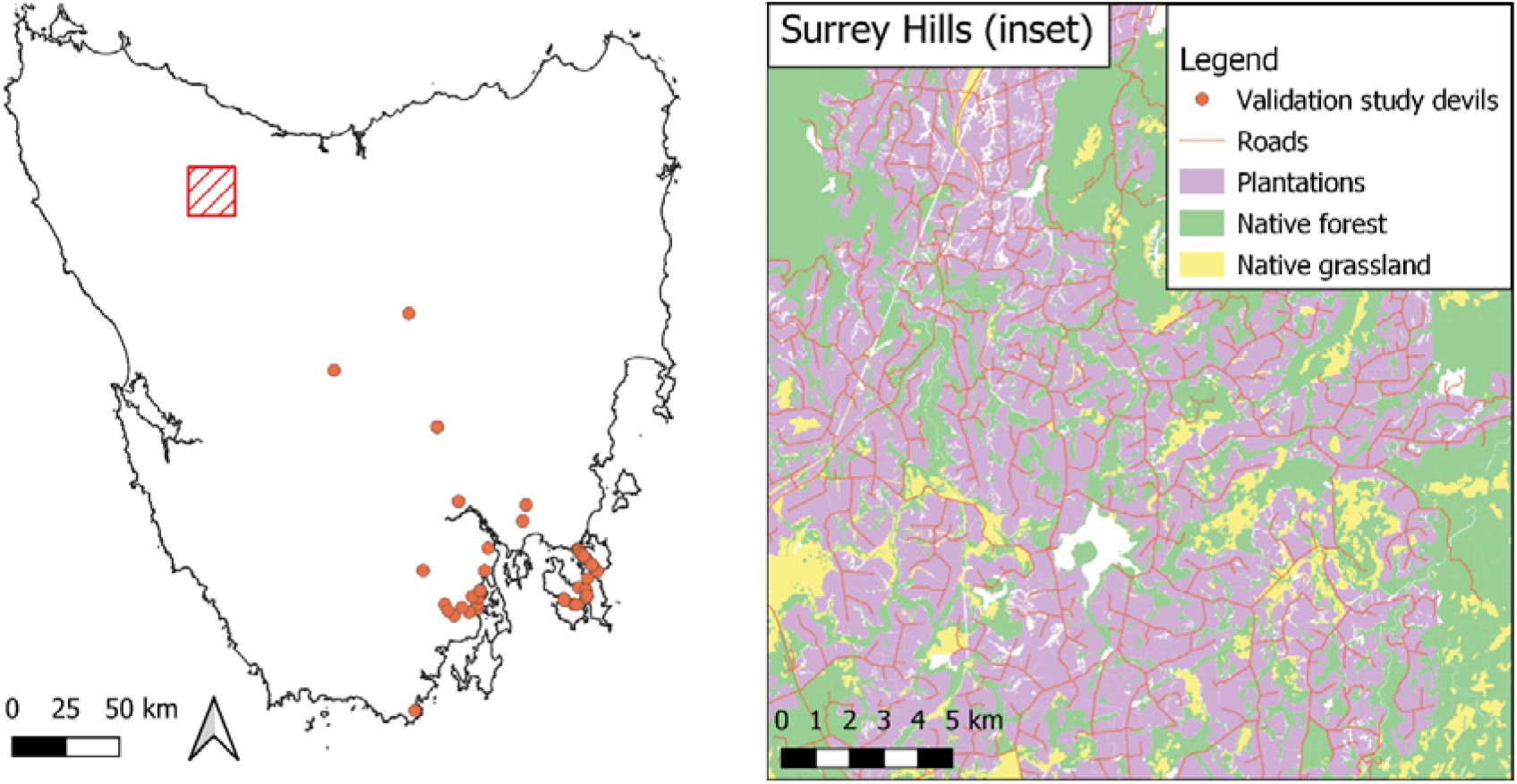
Map of Tasmania, showing where 33 of the 39 Tasmanian devils (*Sarcophilus harrisii*) in the validation study were collected; and Surrey Hills, a eucalypt plantation where all the hair samples for the plantation study were collected. The collection locations of the remaining six devils in the validation study is unknown.

### Forestry plantation landscape study

Relatively small (∼0.1 g) hair samples were collected from 61 live wild devils within Surrey Hills, a forestry plantation landscape in northwest Tasmania (-41.420S, 145.814E) (Figure 1). Surrey Hills is an intensively managed forestry landscape dominated by *Eucalyptus nitens* plantations^28^. Culling, using lead-based ammunition, is regularly undertaken at the site prior to planting in coupes to protect seedlings from browsing by three marsupial species: Bennett’s wallaby, rufous-bellied pademelon, and brushtail possum. All three species are common prey of devils^33^ and there is evidence that culled browsers are scavenged by devils^32^. Culling occurs year-round, and devils are highly reliant on scavenging throughout the year^53^, therefore differences in sampling times are unlikely to have had a major impact on hair lead levels in devils. Between February 2021 and March 2022, we captured devils in custom-made 80 cm by 30 cm PVC pipe traps, shaved a 1 cm^2^ patch of fur from their right shoulders, and recorded the age (from tooth wear) and sex of the animal. These samples were analyzed using XRF only, as they were not large enough for traditional ICP-MS methods. As above, samples were not washed prior to analysis.

### XRF measurements

For XRF analysis of hair samples, we used almost identical methods to those used in past XRF studies of lead exposure in bird bones^14,46,47,52,54^ – see those studies for more detailed descriptions of the process. The portable XRF device used was a Niton XL3t GOLDD+ (Thermo□Fisher Scientific, Omaha, USA), with the same settings, filter combinations and read times as those used to analyze eagle bones in Hampton et al. (2021)^14^. To analyze the spectra produced, we used standard procedures as described in past studies^14,55^.

### ICP-MS measurements

The same hair samples (validation study samples only) were then sent to a commercial laboratory (Edith Cowan University Analytical Facility, Joondalup, Australia) to determine their lead concentrations via inductively coupled plasma mass spectrometry (ICP-MS), as per standard procedure in wildlife toxicology studies. Briefly, hair samples were first freeze-dried, then underwent acid digestion and were homogenized and processed as per liver and bone samples in Pay et al. (2021)^16^. Likewise, lead concentrations from the samples were analyzed with ICP-MS using the same methods as Pay et al. (2021)^16^. We used the same Certified Reference Material (CRM) as Pay et al. (2021)^16^ as a positive control. The CRM was digested twice, and two ICP-MS readings were made on each digestion, producing an average accuracy of 100.2%. We created duplicate reads by re-analyzing every fifth or sixth sample, yielding an average relative standard deviation (RSD) of 2.2%. We also conducted duplicate blind sample digestions on five randomly selected samples (average RSD = 2.0%). The lead concentrations were measured in mg/kg of dry weight (equivalent to ppm). Limit of detection (LOD) was 0.0015 mg/kg and limit of quantification (LOQ) was 0.005 mg/kg. No samples had ICP-MS lead concentrations below the LOD or LOQ.

### Statistical analysis

We identified the uncertainty values for XRF measurements as per past studies^45^. We calculated the uncertainty (*σ*) of each measurement using the following function **(eq 1)**:

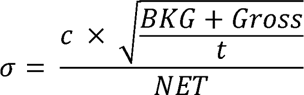

where *c* is concentration, *BKG* is the background counts as estimated by our fitting, Gross is the gross counts, *t* is measurement time, and *NET* is net lead counts from a Gaussian function^14^. The uncertainty values for XRF give the confidence intervals associated with a point estimate of elemental concentration. We retained the negative values for lead in hair because they represent a point estimate of hair lead concentration incorporating uncertainty^56,57^.

We used linear regression to examine the relationship between ICP-MS-derived and XRF-derived lead concentrations for samples tested via both methods (n = 39). We square-root transformed both variables to minimize heteroscedasticity, and estimated the relationship using the following function **(eq 2):**

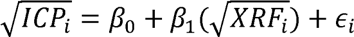

where *β_0_* is the intercept, *β_1_* is the slope of the relationship, and □*_i_* is the residual error. We evaluated predictive competence of the regression model with 10-fold cross-validation, repeated three times. Modelling was conducted using the caret package^58^ in R^59^.

For the plantation study, we used the equation from the regression analysis (Equation 3 below) to predict ICP-MS values from XRF measurements of lead levels in devil hair. We used a t-test to identify any difference in lead levels between devils in the validation and plantation study, and a two-way analysis of variance (ANOVA) to test for differences in lead levels among devils of different sexes and ages in the plantation study (juvenile < 1 year, subadult 1–2 years, adult > 2 years). Additionally, we constructed linear models with the predictors of sex and age to look for any influence of these parameters on predicted ICP-MS lead levels in devils, using Akaike’s Information Criterion (AIC) values to select the best fitting model^60^. Lead level values were log transformed for these analyses, which were undertaken in the R statistical environment^59^.

## RESULTS

We found a positive relationship between lead concentrations in the hair of Tasmanian devils estimated using portable XRF and ICP-MS (n = 39) (Figure 2). The regression model showed a reasonable correlation between portable XRF and ICP-MS values, explaining 61% of the variability in the data over the range of values observed. Expected ICP-MS estimates were predicted from portable XRF estimates across the range of data from the plantation study (n = 61) using the following function (**eq 3**):

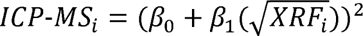

where *β>*_0_ = -0.205 (SE = 0.11) and *β*_1_ = 0.907 (SE = 0.072) (R^2^ = 0.61, t_37_ = 12.651, p < 0.001; RMSE = 0.45).

**Figure 2.**
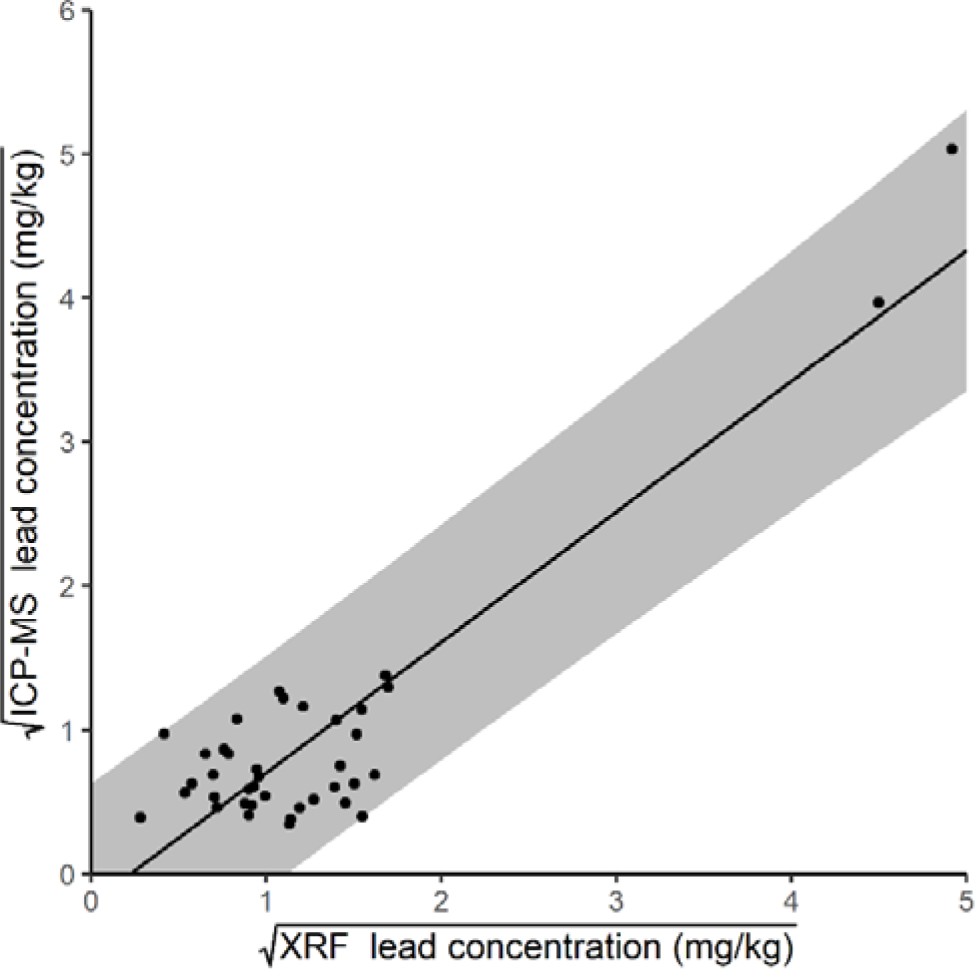
Correlation between portable XRF and ICP-MS lead concentration measurements in Tasmanian devil (*Sarcophilus harrisii*) hair, including all observed values. Shaded polygon represents the 95% prediction interval.

As the two highest values exerted a high degree of influence over the form of the regression, we ran a second model excluding these values. This model showed limited, though still significant, correlation between XRF and ICP-MS values, explaining 43% of the variability in the data (R^2^ = 0.43, t_37_ = 1.869, p < 0.001; RMSE = 0.28).

Most of the samples (94.87% as measured via either method) showed lead concentrations of < 3 mg/kg. However, the two points with the highest lead levels (> 15 mg/kg) measured consistently high with both XRF and ICP-MS (Figure 2). XRF estimates tended to be higher than ICP-MS: mean concentrations were 2.33 mg/kg when measured via XRF versus 1.65 mg/kg when measured via ICP-MS (Table 1).

**Table 1.** Lead Concentration in Hair Samples from 39 Tasmanian Devils (*Sarcophilus Harrisii*), Estimated using Inductively-Coupled Mass Spectrometry (ICP-MS) and Portable X-Ray Fluorescence

|  | Mean | Min | Max | SD |
| --- | --- | --- | --- | --- |
| ICP-MS (mg/kg) | 1.65 | 0.12 | 25.34 | 4.61 |
| Portable XRF (mg/kg) | 2.33 | 0.08 | 25.16 | 4.76 |
| Portable XRF uncertainty (mg/kg) | 0.58 | 0.20 | 1.97 | 0.39 |

Table 2 shows portable XRF hair lead measurements and uncertainties from devils in the plantation landscape. These data were converted to ICP-MS values using equation 3. Two negative values (-1.39 and -0.48 mg/kg, σ = 2.02 and 1.04 respectively) were excluded from analysis as they could not be square-root transformed. There was no detectable difference in mean lead levels between devils in the plantation (mean 1.05 ± 1.02 SD mg/kg) and validation (mean 1.65 ± 4.61) studies (df = 40, t = 0.79, p = 0.43) (Figure 3). In the plantation study, there was no evidence of differences in hair lead levels between sexes (df = 1, F = 0.96, p = 0.33) or among ages (df = 2, F = 0.23, p = 0.79) or an interaction between these two variables (df = 2, F = 1.32, p = 0.28). The best fitting model predicting lead levels in devil hair in this landscape was the null model, indicating no influence of age or sex on lead levels in devils (Table 3). However, these results should be interpreted with caution due to low sample sizes (Figure 4).

**Table 2.** Lead Concentrations in Hair Derived from Portable XRF and Predicted ICP-MS Levels from Regression Analysis for Live Tasmanian Devils (*Sarcophilus Harrisii*), Sampled During a Trapping

|  | Mean | Min | Max | SD |
| --- | --- | --- | --- | --- |
| Predicted ICP-MS (mg/kg) | 1.05 | 0.01 | 4.34 | 1.02 |
| Portable XRF (mg/kg) | 1.69 | -1.39 | 6.37 | 1.53 |
| Portable XRF uncertainty (mg/kg) | 0.85 | 0.25 | 2.02 | 0.45 |

**Table 3.** Model Ranking Testing the Importance of Age and Sex for Predicting Lead Levels in Tasmanian Devil (*Sarcophilus Harrisii*) Hair Samples from a Forestry Plantation Landscape. Df = Degrees of Freedom, AIC = Akaike Information Criterion, ΔAIC = Akaike Information Criterion Difference, AICwt = Akaike Information Criterion Weight

| Model | df | AIC | ΔAIC | AICwt |
| --- | --- | --- | --- | --- |
| Null | 2 | 221.5 | 0 | 0.47 |
| Sex | 3 | 222.8 | 1.24 | 0.25 |
| Age | 4 | 223.5 | 1.96 | 0.18 |
| Age + sex | 5 | 224.7 | 3.19 | 0.10 |

**Figure. 3.**
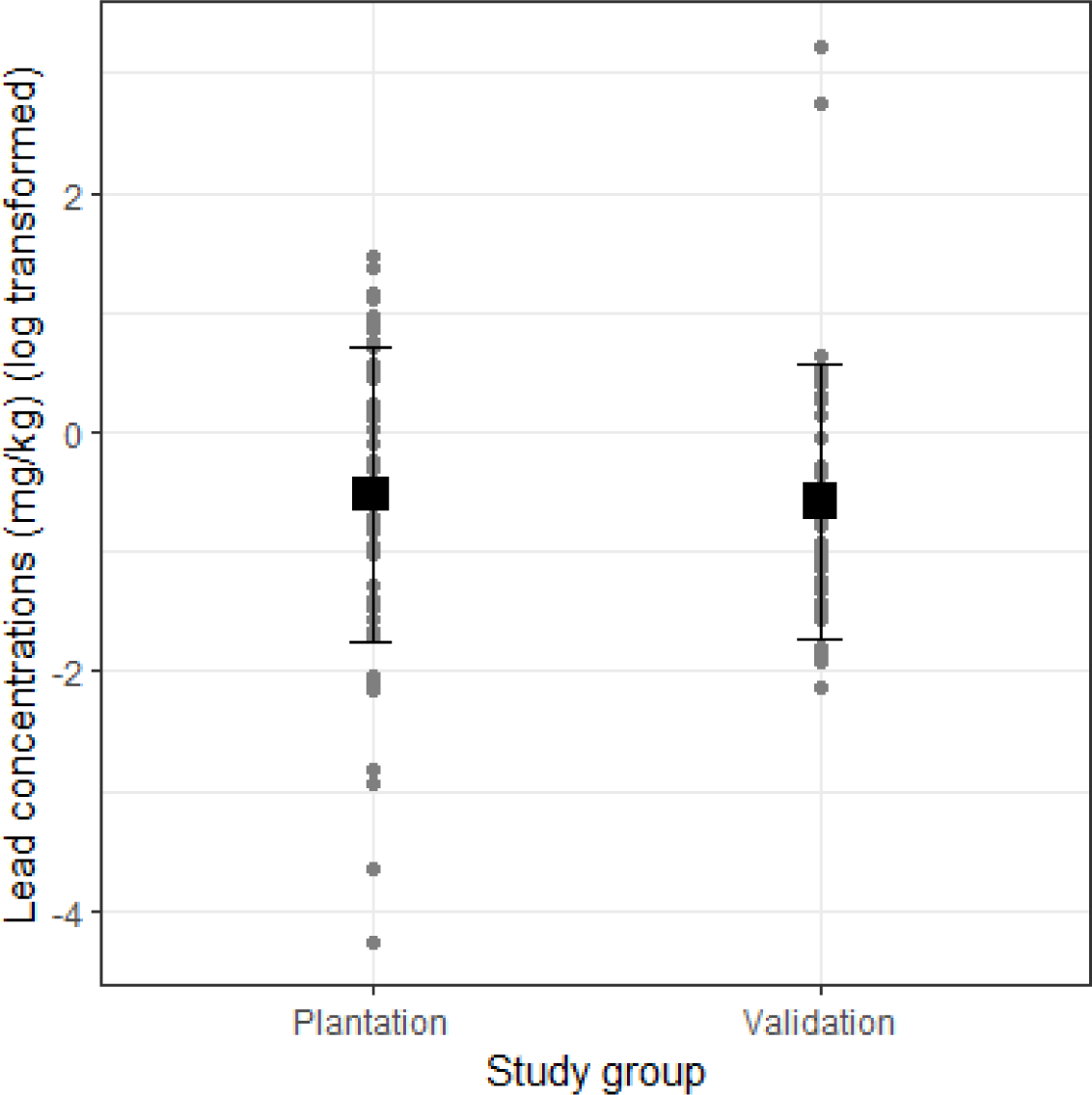
Comparison of lead concentrations in the hair of Tasmanian devils (*Sarcophilus harrisii*) from the plantation study and the validation study, estimated using portable X-ray fluorescence. Points show individual measurements, squares show means and error bars show SDs.

**Figure 4.**
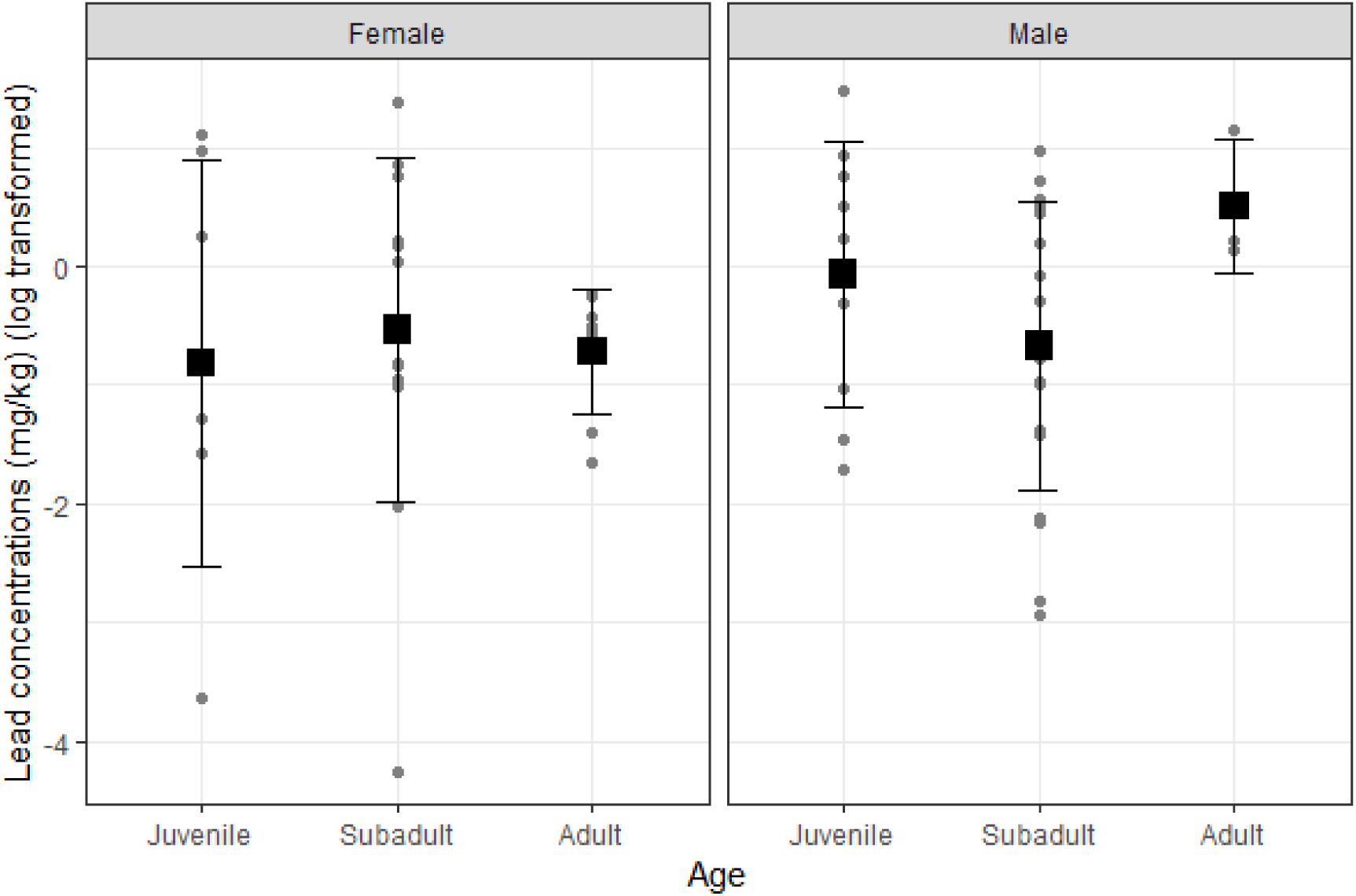
Portable X-ray fluorescence estimates of lead concentration in the hair of Tasmanian devils (*Sarcophilus harrisii*) in a timber plantation landscape. Points show individual measurements, squares show means and error bars show SDs.

## DISCUSSION

We present the first investigation of portable XRF for measurement of elemental concentrations in hair in a non-human species, and the first report of lead exposure in a morphologically specialized mammalian scavenger as measured via hair lead levels. Using regression analysis, we showed reasonable correlation (61%) between XRF and ICPS-MS values. This suggests that portable XRF may be useful as an initial screening method for mammalian hair to identify individuals with the largest burden of lead exposure versus a typical exposure profile, though it is does not seem to be useful for precise predictions of lead exposure levels.

Lead in hair has been found to correlate with lead levels in other tissues such as blood, liver and bone of other species^61,62^. However, this is not always the case; for example, hair lead levels did not correlate with liver lead levels in European hedgehogs (*Erinaceus europaeus*)^62^, or liver and kidney samples in caribou (*Rangifer tarandus caribou*)^63^, though the latter may have been influenced by low lead concentrations. Use of hair and other keratinized matrices to measure lead exposure in mammals should therefore be treated with caution, as small variations in hair lead may not reflect differences in lead concentrations in internal tissues or blood^44^. Given these limitations, XRF may be most useful for distinguishing individuals with substantially elevated hair lead levels from those individuals with background lead levels, rather than precise measurement of hair lead concentrations. Our results indicate portable XRF is likely to be sufficient for this type of ‘screening’ task. Additionally, characterizing the relationship between lead levels in hair and internal tissues/blood of mammalian scavengers such as devils would assist with interpreting lead exposure from hair lead measurements.

Lead levels in devil hair, in both the validation study and the plantation landscape study, were comparable to other published research on lead exposure in mammals (both wild and domestic). The levels we found were lower than those found in the hair of domestic dogs (*Canis familiaris*) used for hunting in Argentina (mean 2.37 ± 6.21 mg/kg)^64^, similar to those in hair from fruit bats in eastern Australia (medians for three species: 1.26/1.64/2.26 mg/kg)^38^ and roe deer in Italy (mean 1.39 ± 1.63 mg/kg)^40^, and higher than those found in hair from Malay civets sampled in Malaysian Borneo (mean 0.52 ± 1.08 mg/kg)^39^. Similar to several of these studies^38–40^, we found no evidence of variation in lead levels in devil hair among sexes and age classes.

Without a standard classification of harmful lead levels in hair, even apparently low levels of lead in devil hair may be cause for concern. Two devils, both in the validation study, showed atypically high hair lead levels: 15.77 and 25.34 mg/kg, when no other devils in either sample group had > 4.4 mg/kg (none > 2 mg/kg in the validation study). The source of these unusually high levels is unknown. The generally low hair lead levels are consistent with other studies showing low lead levels in the blood^25^ and liver^18^ of wild devils. Mean lead levels were similar in devils in the plantation study and the validation study, providing little support for our hypothesis that lead contamination would be higher in devils living in a modified landscape subject to regular culling.

Grey wolves (*Canis lupus*), cougars (*Puma concolor*) and American black bears (*Ursus americanus*) in the greater Yellowstone ecosystem likewise showed relatively low lead levels in blood and tissue samples, indicating that some mammalian scavengers in some areas may not be impacted by lead exposure even where hunting using lead-based ammunition occurs^34^. Brown bears did show elevated lead levels in this landscape, but lead levels did not increase during the hunting season^34^. In contrast, brown bears sampled in northern Europe have shown elevated lead levels in blood and milk^21^, which in one study was linked to hunting activity^22^, and lead levels in the teeth of Canadian black bears increase markedly during hunting seasons^23^. It is clear that further research is required to identify risk factors and determine harmful lead exposure in free ranging mammalian species^20^.

One limitation to the results of this study is that we did not wash the samples in this study prior to analysis, as we aimed to validate portable XRF for use in the field, and washing the fur of live-caught animals would be impractical. Samples may therefore have been contaminated by external lead^41,65,66^. However, lead levels in devil hair in our study were generally low despite the possibility of external contamination. Nonetheless, we could not distinguish between endogenous lead incorporated into the hair from the blood, and exogenous lead from external contamination, in our XRF measurements. We recommend further research to corroborate our results by comparing ICP-MS and portable XRF values of washed hair samples. Another limitation relates to operator safety. The handheld XRF emits a small amount of radiation, which for standard use would be indistinguishable from background radiation in terms of dosimetry^67^. For those planning on regular use of these systems in the field, proper dosimetry should be used to monitor any potential exposure to the user.

In conclusion, Portable XRF has the potential to be developed into a non-destructive, non-invasive method of analyzing lead levels in mammal hair. We found sufficient correlation between lead levels in devil hair derived via portable XRF and ICP-MS to identify differences in lead concentrations over a coarse scale. Our preliminary data suggest portable XRF for hair lead analysis has promise as a screening test to distinguish animals with marked lead exposure from those with relatively low lead levels. While ICP-MS remains the more accurate and precise method to quantify lead exposure levels in non-human mammalian hair, portable XRF is a non-invasive alternative in situations where invasive sample collection or destructive analysis is impractical or nonpreferred. These contexts may include when results are required immediately, for use in the field, when working with threatened species, or for archived museum specimens. Notably, any attempts to use portable XRF as a field-based tool for measuring hair lead concentrations in live animals would need to be cognizant of health and safety risks and regulations associated with the use of an ionizing radiation source^51^. While radiation exposure to the user can be minimized with training and proper use, it is a consideration when choosing a method for lead monitoring. Future research could test the relationship between ICP-MS and portable XRF hair lead measurements in other mammalian scavenging species across a larger lead concentration gradient to refine this method.

## ETHICS AND PERMITS

The live animal trapping and sample collection component of this study was approved by the University of Tasmania Animal Ethics Committee (AEC), approval number 23211, and conducted under a permit to take threatened fauna for scientific purposes (#TFA 21284) from the Tasmanian Department of Natural Resources and Environment. Sample collection from dead animals was performed under permit #TFA 21284. Samples were shipped interstate for laboratory analysis under a Wildlife Export Permit (#TFA 21284) from the Tasmanian Department of Natural Resources and Environment.

## ACKNOWLEDGEMENTS

We acknowledge the Palawa people as the Traditional Owners of the lands on which this research was conducted. For access to samples, we acknowledge the following people: Bill Brown and Sarah Peck from the Department of Natural Resources and Environment Tasmania, and David Hocking and Nicole Zehntner from the Tasmanian Museum and Art Gallery (TMAG). For assistance with dissections, we thank Regi Broeren. For laboratory analysis, we thank Mark Bannister from Edith Cowan University. For fauna licenses needed to import samples to Victoria, we thank Sue Hadden and Roberta Campbell from the Victorian Department of Environment, Land and Water. For facilitating XRF hire, we thank Shellby Thomas-Brookes, Denise Guldemond and Alex Toy from Portable Analytical Solutions. We thank Forico Pty Ltd for providing accommodation and access to land they manage for fieldwork.

This work was supported by the Paddy Pallin Foundation [grant number H0029051], the McKenzie Fellowship Program of the University of Melbourne [grant number 000376], the Holsworth Wildlife Research Endowment [Grant No. H0027981], the Dr Eric Guiler Tasmanian Devil Research Grant, and a matched funding scheme through Forests and Wood Products Australia Limited, with funding provided by the Forest Practices Authority, Forico Pty Ltd, SFM, Sustainable Timber Tasmania, Timberlands, Norske Skog, and Private Forests Tasmania. Dr. Aaron Specht was partially funded by a training grant from the National Institute of Occupational Safety and Health [grant number K01OH012528].

